# Durability of immune responses to the BNT162b2 mRNA vaccine

**DOI:** 10.1101/2021.09.30.462488

**Authors:** Mehul S. Suthar, Prabhu S. Arunachalam, Mengyun Hu, Noah Reis, Meera Trisal, Olivia Raeber, Sharon Chinthrajah, Meredith E. Davis-Gardner, Kelly Manning, Prakriti Mudvari, Eli Boritz, Sucheta Godbole, Amy R. Henry, Daniel C. Douek, Peter Halfmann, Yoshihiro Kawaoka, Veronika I. Zarnitsyna, Kari Nadeau, Bali Pulendran

**Affiliations:** Emory University School of Medicine, Atlanta, GA, USA; Institute for Immunity, Transplantation and Infection, Stanford University, Stanford, CA, USA; Department of Medicine, Division of Pulmonary, Allergy and Critical Care Medicine, Stanford, CA, USA; Vaccine Research Center, National Institute of Allergy and Infectious Diseases, National Institutes of Health, Bethesda, MD, USA; University of Wisconsin, Madison, WI, USA; National Center for Global Health and Medicine, Tokyo, Japan; University of Tokyo, Tokyo, Japan; Department of Pediatrics, Emory Vaccine Center, Yerkes National Primate Research Center, Emory University School of Medicine, Atlanta, GA, USA; Sean N. Parker Center for Allergy & Asthma Research, Stanford, CA, USA; Department of Pathology, Stanford University School of Medicine, Stanford University, Stanford, CA, USA; Department of Microbiology and Immunology, Stanford University School of Medicine, Stanford University, Stanford, CA, USA

## Abstract

The development of the highly efficacious mRNA vaccines in less than a year since the emergence of SARS-CoV-2 represents a landmark in vaccinology. However, reports of waning vaccine efficacy, coupled with the emergence of variants of concern that are resistant to antibody neutralization, have raised concerns about the potential lack of durability of immunity to vaccination. We recently reported findings from a comprehensive analysis of innate and adaptive immune responses in 56 healthy volunteers who received two doses of the BNT162b2 vaccination. Here, we analyzed antibody responses to the homologous Wu strain as well as several variants of concern, including the emerging Mu (B.1.621) variant, and T cell responses in a subset of these volunteers at six months (day 210 post-primary vaccination) after the second dose. Our data demonstrate a substantial waning of antibody responses and T cell immunity to SARS-CoV-2 and its variants, at 6 months following the second immunization with the BNT162b2 vaccine. Notably, a significant proportion of vaccinees have neutralizing titers below the detection limit, and suggest a 3^rd^ booster immunization might be warranted to enhance the antibody titers and T cell responses.

---

The development of the highly efficacious mRNA vaccines in less than a year since the emergence of SARS-CoV-2 represents a landmark in vaccinology. The efficacy of the Pfizer-BioNTech (BNT162b2) and Moderna (mRNA1273) vaccines were 95% (95% confidence limits 90.3% to 97.6%)^1^ and 94.1% (95% confidence limits 89.3% to 96.8%)^2^, respectively, for a median of 2 months following the second dose. However, reports of waning vaccine efficacy^3-5^, coupled with the emergence of variants of concern that are resistant to antibody neutralization, have raised concerns about the potential lack of durability of immunity to vaccination. While the durability of antibody responses to the mRNA1273 vaccine has been reported recently^5^, there is a lack of data on the durability of immune responses induced by the BNT162b2. We recently reported findings from a comprehensive analysis of innate and adaptive immune responses in 56 healthy volunteers who received two doses of the BNT162b2 vaccination^6^. Here, we analyzed antibody responses to the homologous Wu strain as well as several variants of concern, including the emerging Mu (B.1.621) variant, and T cell responses in a subset of these volunteers at six months (day 210 post-primary vaccination) after the second dose.

Of the 56 volunteers, we were able to collect blood samples from 46 participants on day 210. Anti-Spike binding IgG response measured in all 46 participants declined 7.3-fold between days 42 and 210, with two individuals showing a response below the limit of detection (**Fig. 1a**). The decline in response was 6.3-fold in female participants and 9.1-fold in male participants, but the difference in responses was not statistically significant (**Fig. 1a**). Consistent with this, the authentic live virus neutralization antibody titers against the homologous USA-WA1/2020 strain were 7-fold lower at 6 months relative to the titers at 3 weeks (**Fig. 1b**). The decline in response was 6.6- and 7.4-folds in female and male participants, respectively (**Fig. 1b**). Of note, we performed this assay in all 46 volunteers in Vero-E6 cells to compare with the data we reported recently^6^ and all the previous studies where we used this assay^5,7-9^. The durability of neutralizing titers against the homologous strain was comparable to that of the responses in mRNA1273 vaccinees^5,10^. To examine this methodically, we calculated decay rates of binding and neutralizing antibody responses using an exponential decay model, which assumes a steady decay rate over time. The estimated half-lives of binding and neutralizing antibody responses were 58 (95% Cl: 53.5 to 62.5) and 56 (95% Cl: 51 to 62) days, respectively (**Fig. 1c**). We also employed the power law model, which assumes a decreased decay rate over time. While the estimated decay rate was 81 days using this model, a corrected Bayesian information criterion used to compare models suggested that the exponential decay model provided the best fit. In comparison, the half-life of live-virus neutralizing antibody response to mRNA1273 vaccination was 68 days (95% Cl: 61 to 75) as described in a previous study^10^. In contrast, the half-life of live-neutralizing antibody response was estimated as 150 days in COVID-19 infected individuals^11^. In the case of other inactivated vaccines such as the diphtheria or tetanus vaccines, the decay rates of the antibody response have been calculated to be in the order of a year or so^12^. In contrast, analysis of decay rates of immune responses induced by live viral vaccines such as those against measles, rubella, vaccinia, mumps, and VZV, performed using the exponential decay modes, indicate essentially no decay in the antibody response, highlighting the remarkable durability of the response stimulated by these vaccines^12^.

**Fig. 1.**
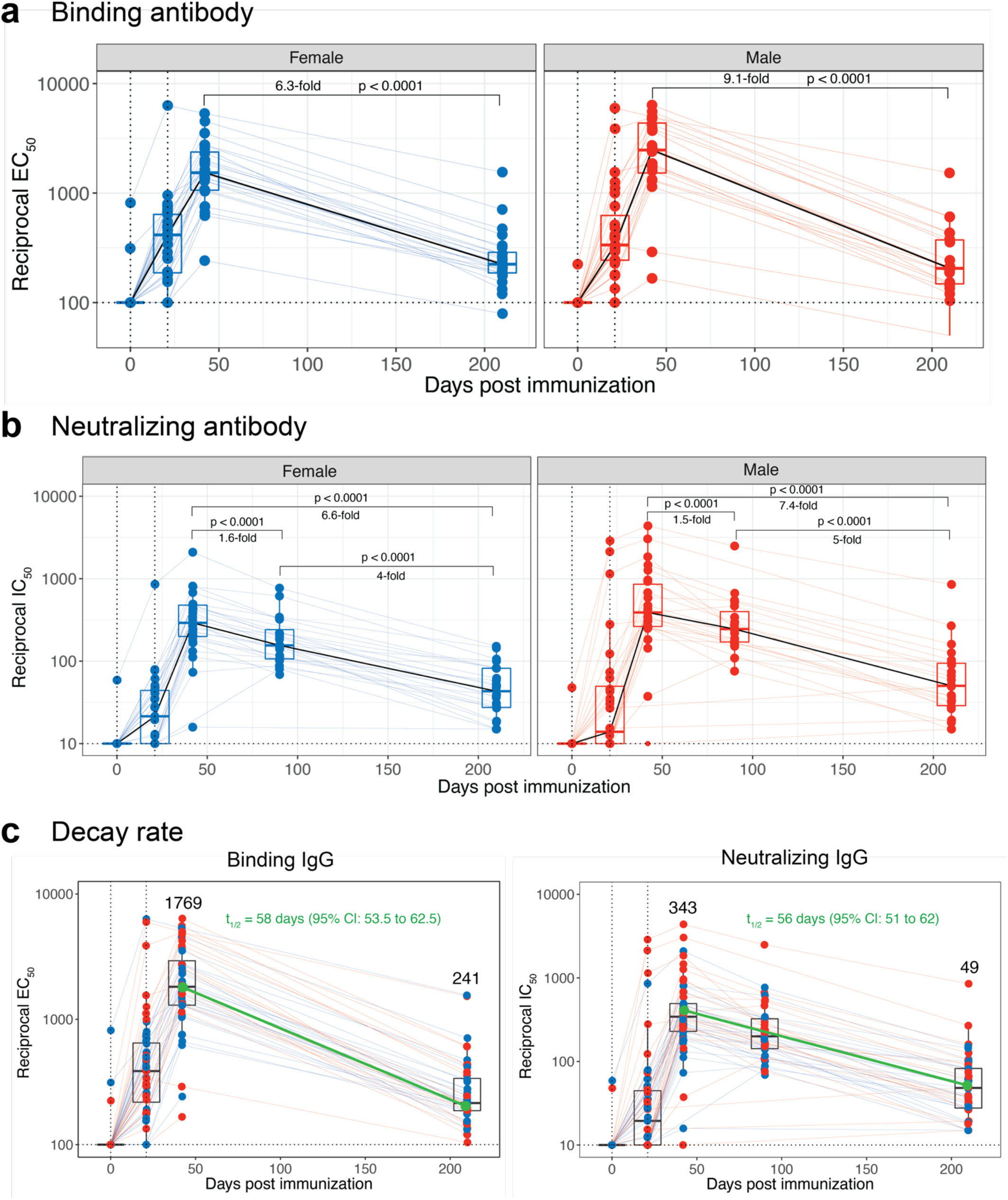
Durability of antibody following the Pfizer-BioNTech mRNA vaccination. **a**, Kinetics of anti-Spike binding IgG titers measured using ELISA (N = 46, 24 females and 22 males on day 210). Data of day 0 and 21 were obtained from our previously published study^6^. Day 42 samples were re-assayed with day 210 samples by ELISA. **b**, Kinetics of authentic live virus neutralizing antibody response against the homologous USA/WA1 strain (N = 46, 24 females and 22 males on day 210). Data of day 0, 21 and 90 were obtained from our previously published study^6^. Day 42 samples were re-assayed with day 210 samples by the FRNT assay. **c**, Kinetics of binding (left) and neutralizing (right) antibody responses of all participants. The dark green line indicates half-life estimated using the exponential decay model. In all the panels, each colored circle represents an individual. Red and blue circles indicate males and females, respectively. All statistical comparisons were non-parametric pair-wise comparisons performed using Wilcoxon’s matched-pairs signed rank test.

Next, we measured T cell responses in 12 individuals to measure the durability of memory T cells. Spike-specific CD4 T cells expressing IFN-γ and TNF were significantly downregulated on day 210 compared to day 28; however, the IL-2^+^ CD4 T cell response did not differ significantly (**Fig. 2a**). The frequency of spike-specific circulating Tfh-like (follicular T helper) cells expressing IL-21 and CD154, critical for their role in the germinal center formation and generation of durable antibody responses, were also significantly downregulated by day 210 (Fig. 2b). However, even at the peak, the frequency was only modest compared to effector cytokines such as IFN-γ and TNF^6^. Similarly, frequencies of antigen-specific CD8 T cells secreting IL-2, IFN-γ, and TNF demonstrated a marked diminution and were below the detection limit in most participants (**Fig. 2c**) on day 210.

**Fig. 2.**
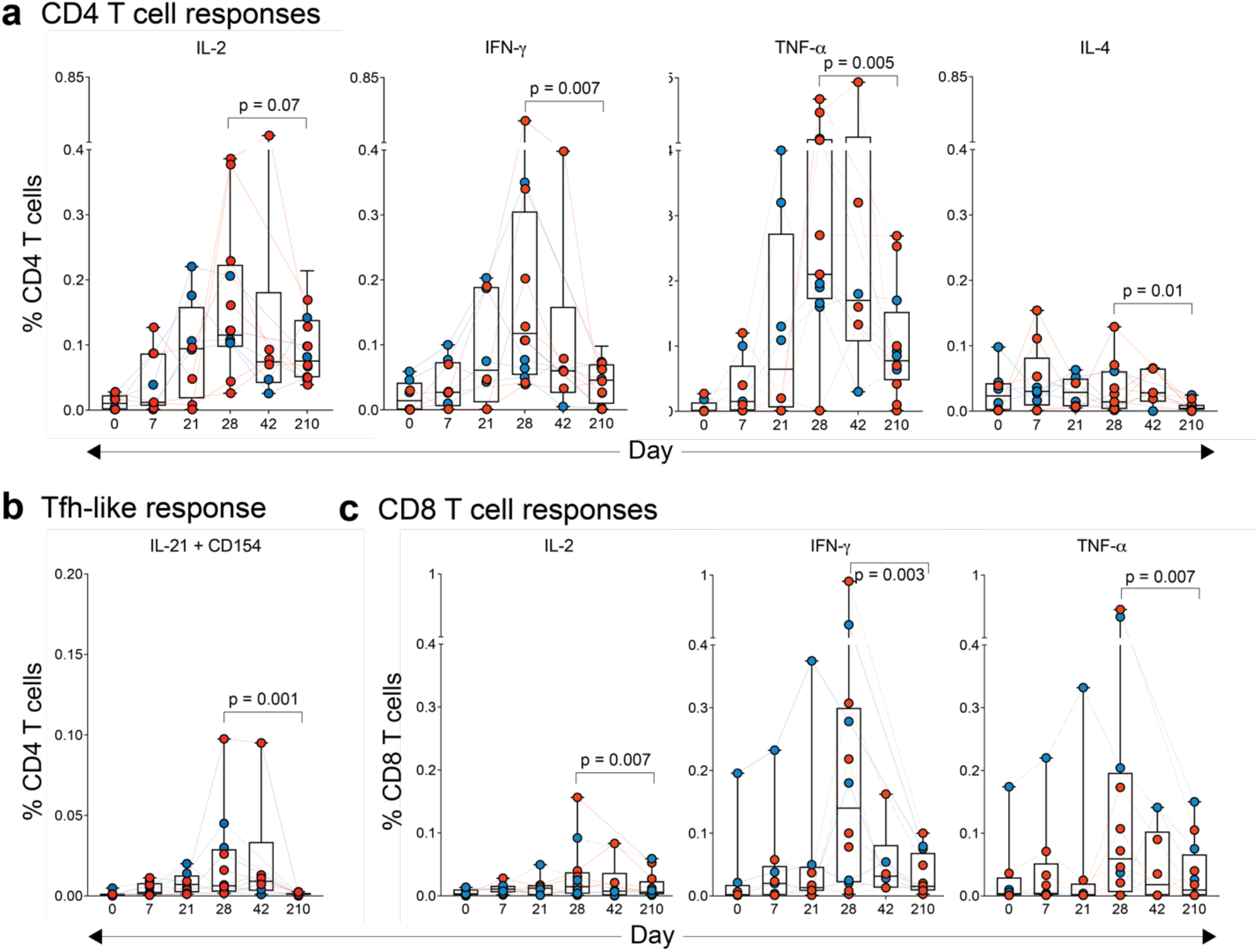
Durability of T cell responses following the Pfizer-BioNTech mRNA vaccination. **a - c**, Frequency of Spike-specific CD4 T cells expressing IL-2, IFN-γ, TNF and IL-4 (**a**); CD4 T cells expressing IL-21 and CD154 (**b**); and CD8 T cells expressing IL-2, IFN-γ and TNF (**c**). In all the panels, data of day 0, 7, 21 and 42 were obtained from our previously published study^6^. Day 28 samples were re-assayed with day 210 samples. Each colored circle represents an individual. Red and blue circles indicate males and females, respectively. All statistical comparisons were non-parametric pair-wise comparisons performed using Wilcoxon’s matched-pairs signed rank test.

The emergence of several variants of concern since the development of BNT162b2 has raised concerns on the cross-neutralization potential of vaccine-induced antibody responses. To evaluate cross-neutralization, we performed an *in vitro* live virus Focus Reduction Neutralization Test (FRNT)^13^ against the Beta (B.1.351), Delta (B.1.617.2), Gamma (P.1), and the recently described Mu (B.1.621) variants in 17 participants. Of note, we used Vero-TMPRSS cells for this assay differing from the data reported in Fig. 1. The FRNT50 GMTs at 3 weeks post-secondary vaccination were 302 for WA1/2020 (95% CI, 181 to 428), 59 for Beta (95% CI, 39 to 99), 175 for Delta (95% CI, 112 to 278), 155 for Gamma (95% CI, 77 to 297), and 130 for Mu (95% CI, 72 to 192) (**Fig. 3a**). There was a significant reduction (up to 7-fold) in the titers against these variants at 6 months. At 6 months, the FRNT50 GMTs were 43 for WA1/2020 (95% CI, 21 to 77), 26 for Beta (95% CI, 15 to 41), 25 for Delta (95% CI, 15 to 35), 44 for Gamma (95% CI, 15 to 77), and 32 for Mu (95% CI, 25 to 102). Furthermore, we compared the efficiency of cross-neutralization of the variants of concern in serum at day 42 and day 210 (**Fig. 3b**). The Beta (B.1.351) variant demonstrated the highest decline compared to the homologous WA1 strain on day 42 (5.2-fold), consistent with previous reports suggesting this as the most difficult-to-neutralize strain owing to mutations associated with immune escape from infection- and vaccine-induced antibodies^14,15^ (**Fig. 3b, left panel**). On day 210, the difference between WA1 and Beta strains was only 1.7-fold, suggesting that potent antibodies that could cross-neutralize were maintained. However, 9 of the 17 participants (53%) had a titer against the Beta variant that were below the detection limit of the assay (**Fig. 3b, right panel**). On the other hand, the neutralizing antibody response to the delta variant (B.1.617.2) that became the dominant circulating strain in several countries worldwide since its first description in October 2020 was reduced 1.7-fold at both day 42 and 210-time points (**Fig. 3b**). Similar to the beta variant, 47% of the volunteers had an undetectable neutralizing response to the delta variant (**Fig. 3b, right panel**). Importantly, we also measured the neutralizing response against the Mu (B.1.621) variant, first reported in Colombia earlier this year, and has recently been added to the growing list of variants of concern. Of note, the response to the Mu variant was 2.3-fold lower in comparison to the WA1 strain in day 42 sera (**Fig. 3b, left panel**). The magnitude of the decline was intermediate between delta and beta variants. On day 210, 53% of participants had titers below the detection limit compared to the beta variant (**Fig. 3b, right panel**).

**Fig. 3.**
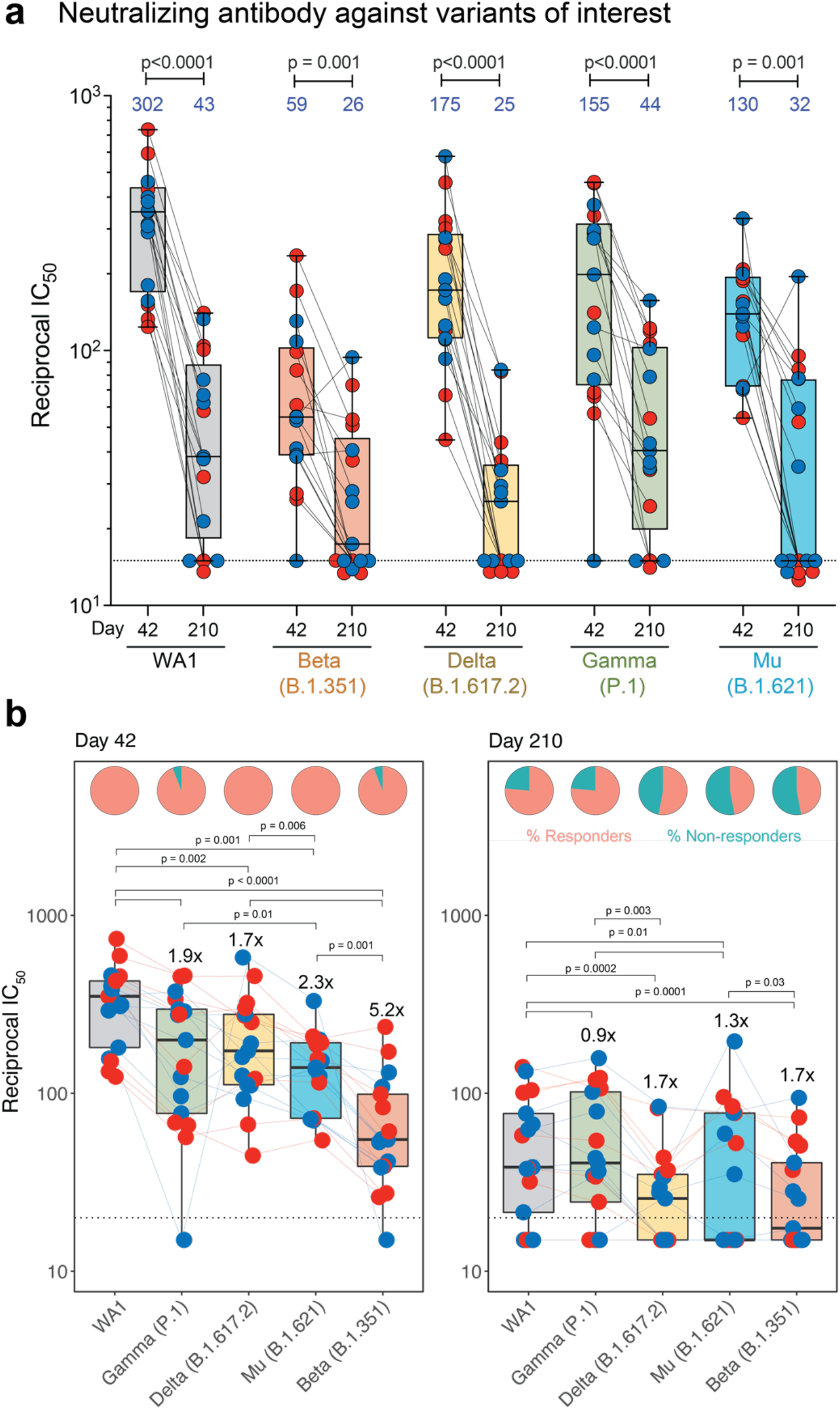
Durability of cross-neutralizing antibody responses following the Pfizer-BioNTech mRNA vaccination. **a**, Authentic live virus neutralizing antibody responses against the homologous USA/WA1 strain and the variants of concerns B.1.351 (Beta), B.1.617.2 (Delta), P.1 (Gamma) and B.1.621 (Mu) (N=17). The numbers in blue indicate geometric mean titers. **b**, Authentic live virus neutralizing antibody responses against the homologous USA/WA1 strain and the variants of concerns B.1.351 (Beta), B.1.617.2 (Delta), P.1 (Gamma) and B.1.621 (Mu) on day 42 (left panel) and day 210 (right panel). The pie charts on top of the bars represent proportion of responders in orange and non-responders in green. In all the panels, each colored circle represents an individual. Red and blue circles indicate males and females, respectively. All statistical comparisons were non-parametric pair-wise comparisons performed using Wilcoxon’s matched-pairs signed rank test.

In summary, these data demonstrate a substantial waning of antibody responses and T cell immunity to SARS-CoV-2 and its variants, at 6 months following the second immunization with the BNT162b2 vaccine. Notably, a significant proportion of vaccinees have neutralizing titers below the detection limit, and suggest a 3^rd^ booster immunization might be warranted to enhance the antibody titers and T cell responses.

## Methods

### Human subjects and experimentation

Fifty-six healthy volunteers were recruited for the study at the Stanford University under informed consent. The study was approved by Stanford University under Institutional Review Board protocols 8269 and 60171. The samples were collected as a part of registered clinical trials NCT04828603 and NCT04664309. The study was conducted within full compliance of Good Clinical Practice as per the Code of Federal Regulations. The demographics of the study participants have been published previously^6^.

### Anti-spike binding enzyme-linked immunosorbent assay

SARS-CoV-2 spike protein was purchased from Sino Biologicals. Ninety-six-well high binding plates were coated with 100 ng of spike protein diluted at a concentration of 2 μg ml^−1^ in PBS. The next morning, the plates were washed once, blocked with 3% non-fat milk in PBS containing 0.1% Tween 20 (PBST) for 1 h at room temperature. Sera samples serially diluted in 1% non-fat milk containing PBST was added to the plates and incubated at 37 °C for 1 h. The plates were washed 3× with PBST, horseradish peroxidase conjugated goat anti-monkey IgG (γ-chain specific, Alpha Diagnostics, 1:4,000 dilution), in PBS-T containing 1% non-fat milk was added and incubated for 1 h at room temperature. Wells were washed 3× with PBST before addition of 3,3′,5,5′-tetramethylbenzidine (TMB) substrate solution. The reaction was stopped after 12 min by addition of 0.16 M sulfuric or 1 M hydrochloric acid. The optical density at 450 nanometres was measured with a Biorad microplate reader.

### Viruses and cells

VeroE6-TMPRSS2 cells were described previously^16^ and cultured in complete DMEM in the presence of Gibco Puromycin 10mg/mL (# A11138-03). nCoV/USA_WA1/2020 (WA/1), closely resembling the original Wuhan strain and resembles the spike used in the mRNA-1273 and Pfizer BioNTech vaccine, was propagated from an infectious SARS-CoV-2 clone as previously described^17^. icSARS-CoV-2 was passaged once to generate a working stock. The hCoV-19/USA/MD-HP01542/2021 (B.1.351) was provided by Dr. Andy Pekosz (John Hopkins University, Baltimore, MD) and propagated in Vero-TMPRSS2 cells^5^. The hCoV-19/Japan/TY7-503/2021 (P.1) was provided by BEI resources and propagated in Vero-TMPRSS2 cells. hCoV-19/USA/WI-UW-4340/2021 was isolated from a residual nasopharyngeal swab sample collected from a health care worker at University Hospital (Madison, WI) in April 2021 as part of a study approved by the institutional review board at the University of Wisconsin (2020-0897). hCoV-19/USA/CA-Stanford-15_S02/2021 (herein referred to as the B.1.617.1 variant) was derived from a mid-turbinate nasal swab collected from a Stanford Healthcare patient in March 2021 and isolated as previously described^16^.

### Virus stock sequencing

All viral working stocks were deep sequenced. Illumina-ready libraries were generated using NEBNext Ultra II RNA Prep reagents (New England BioLabs) as previously described^18^. Briefly, we fragmented RNA, followed by double-stranded cDNA synthesis, end repair, and adapter ligation. The ligated DNA was then barcoded and amplified by a limited cycle PCR and the barcoded Illumina libraries were sequenced by using paired-end 150-base protocol on a NextSeq 2000 (Illumina). Demultiplexed sequence reads were analyzed in the CLC Genomics Workbench v.21.0.3 by (i) trimming for quality, length, and adaptor sequence, (ii) mapping to the Wuhan-Hu-1 SARS-CoV-2 reference (GenBank accession number: NC_045512), (iii) improving the mapping by local realignment in areas containing insertions and deletions (indels), and (iv) generating both a sample consensus sequence and a list of variants. Default settings were used for all tools.

### Focus Reduction Neutralization Test

FRNT assays were performed as previously described^13^. Briefly, samples were diluted at 3-fold in 8 serial dilutions using DMEM (VWR, #45000-304) in duplicates with an initial dilution of 1:10 in a total volume of 60 μl. Serially diluted samples were incubated with an equal volume of WA1/2020 or B.1.617.1 or B.1.617.2 (100-200 foci per well based on the target cell) at 37° C for 1 hour in a round-bottomed 96-well culture plate. The antibody-virus mixture was then added to VeroE6-TMPRSS2 cells and incubated at 37° C for 1 hour. Post-incubation, the antibody-virus mixture was removed and 100 μl of pre-warmed 0.85% methylcellulose (Sigma-Aldrich, #M0512-250G) overlay was added to each well. Plates were incubated at 37° C for 16 hours. After 16 hours, methylcellulose overlay was removed, and cells were washed three times with PBS. Cells were then fixed with 2% paraformaldehyde in PBS for 30 minutes. Following fixation, plates were washed twice with PBS and 100 μl of permeabilization buffer, was added to the fixed cells for 20 minutes. Cells were incubated with an anti-SARS-CoV spike primary antibody directly conjugated with alexaflour-647 (CR3022-AF647) for up to 4 hours at room temperature. Cells were washed three times in PBS and foci were visualized and imaged on an ELISPOT reader (CTL).

### Quantification and Statistical Analysis

Antibody neutralization was quantified by counting the number of foci for each sample using the Viridot program^19^. The neutralization titers were calculated as follows: 1- (ratio of the mean number of foci in the presence of sera and foci at the highest dilution of respective sera sample). Each specimen was tested in duplicate. The FRNT-50 titers were interpolated using a 4-parameter nonlinear regression in GraphPad Prism 8.4.3. Samples that do not neutralize at the limit of detection at 50% are plotted at 15 and was used for geometric mean calculations.

### T cell intracellular cytokine assay

Live frozen PBMCs were revived, counted, and resuspended at a density of 2 million live cells per ml in complete RPMI (RPMI supplemented with 10% FBS and antibiotics). The cells were rested overnight at 37°C in a CO_2_ incubator. The next morning, the cells were counted again, resuspended at a density of 15 million cells per ml in complete RPMI and 100 μl of cell suspension containing 1.5 million cells was added to each well of a 96-well round-bottomed tissue culture plate. Each sample was treated with two conditions, no stimulation and a peptide pool spanning the spike protein at a concentration of 1 μg ml^−1^ of each peptide in the presence of 1 μg ml^−1^ of anti-CD28 (clone CD28.2, BD Biosciences) and anti-CD49d (clone 9F10, BD Biosciences) as well as anti-CXCR3 and anti-CXCR5. The peptides were custom synthesized to 90% purity using GenScript, a commercial vendor. All samples contained 0.5% v/v DMSO in total volume of 200 μl per well. The samples were incubated at 37°C in CO_2_ incubators for 2 h before addition of 10 μg ml^−1^ Brefeldin-A. The cells were incubated for an additional 4 h. The cells were washed with PBS and stained with Zombie UV fixable viability dye (Biolegend). The cells were washed with PBS containing 5% FCS, before the addition of surface antibody cocktail. The cells were stained for 20 min at 4 °C in 100 μl volume. Subsequently, the cells were washed, fixed and permeabilized with cytofix/cytoperm buffer (BD Biosciences) for 20 min. The permeabilized cells were stained with intracellular cytokine staining antibodies for 20 min at room temperature in 1X perm/wash buffer (BD Biosciences). The details of antibodies are reported in the reporting summary. Cells were then washed twice with perm/wash buffer and once with staining buffer before acquisition using the BD Symphony Flow Cytometer and the associated BD FACS Diva software. All flow cytometry data were analysed using Flowjo software v.10 (TreeStar).

## Acknowledgments

This work was supported in part by NIH grants HIPC U19AI090023 (to B.P.), U19AI057266 (to B.P. and principal investigator R. Ahmed (Emory University)); Open Philanthropy (to B.P.); the Sean Parker Cancer Institute; the Soffer endowment (to B.P.); the Violetta Horton endowment (to B.P.); NIH P51 OD011132, 1U54CA260563 and HHSN272201400004C to Emory University and 75N93021C00014 to the University of Wisconsin from the National Institute of Allergy and Infectious Diseases (NIAID), National Institutes of Health (NIH), by intramural funding from the National Institute of Allergy and Infectious Diseases, Emory Executive Vice President for Health Affairs Synergy Fund award, the Pediatric Research Alliance Center for Childhood Infections and Vaccines and Children’s Healthcare of Atlanta, the Emory-UGA Center of Excellence for Influenza Research and Surveillance (Atlanta, GA USA), COVID-Catalyst-I^3^ Funds from the Woodruff Health Sciences Center and Emory School of Medicine, Woodruff Health Sciences Center 2020 COVID-19 CURE Award; the Parker foundation (to K.C.N.); and the Crown foundation (to K.C.N.). Funders played no role in the design and conduct of the study; collection, management, analysis, and interpretation of the data; preparation, review, or approval of the manuscript; and decision to submit the manuscript for publication.

## References

1 Polack, F. P. et al. Safety and Efficacy of the BNT162b2 mRNA Covid-19 Vaccine. N Engl J Med 383, 2603–2615, doi:10.1056/NEJMoa2034577 (2020).

2 Baden, L. R. et al. Efficacy and Safety of the mRNA-1273 SARS-CoV-2 Vaccine. N Engl J Med 384, 403–416, doi:10.1056/NEJMoa2035389 (2021).

3 Thomas, S. J. et al. Safety and Efficacy of the BNT162b2 mRNA Covid-19 Vaccine through 6 Months. N Engl J Med, doi:10.1056/NEJMoa2110345 (2021).

4 Goldberg, Y. et al. Waning immunity of the BNT162b2 vaccine: A nationwide study from Israel. medRxiv, doi:https://doi.org/10.1101/2021.08.24.21262423 (2021).

5 Pegu, A. et al. Durability of mRNA-1273 vaccine-induced antibodies against SARS-CoV-2 variants. Science, doi:10.1126/science.abj4176 (2021).

6 Arunachalam, P. S. et al. Systems vaccinology of the BNT162b2 mRNA vaccine in humans. Nature, doi:10.1038/s41586-021-03791-x (2021).

7 Anderson, E. J. et al. Safety and Immunogenicity of SARS-CoV-2 mRNA-1273 Vaccine in Older Adults. N Engl J Med, doi:10.1056/NEJMoa2028436 (2020).

8 Suthar, M. S. et al. Rapid Generation of Neutralizing Antibody Responses in COVID-19 Patients. Cell Rep Med 1, 100040, doi:10.1016/j.xcrm.2020.100040 (2020).

9 Widge, A. T. et al. Durability of Responses after SARS-CoV-2 mRNA-1273 Vaccination. N Engl J Med, doi:10.1056/NEJMc2032195 (2020).

10 Doria-Rose, N. et al. Antibody Persistence through 6 Months after the Second Dose of mRNA-1273 Vaccine for Covid-19. N Engl J Med, doi:10.1056/NEJMc2103916 (2021).

11 Cohen, K. W. et al. Longitudinal analysis shows durable and broad immune memory after SARS-CoV-2 infection with persisting antibody responses and memory B and T cells. Cell Rep Med 2, 100354, doi:10.1016/j.xcrm.2021.100354 (2021).

12 Antia, A. et al. Heterogeneity and longevity of antibody memory to viruses and vaccines. PLoS Biol 16, e2006601, doi:10.1371/journal.pbio.2006601 (2018).

13 Vanderheiden, A. et al. Development of a Rapid Focus Reduction Neutralization Test Assay for Measuring SARS-CoV-2 Neutralizing Antibodies. Curr Protoc Immunol 131, e116, doi:10.1002/cpim.116 (2020).

14 Wibmer, C. K. et al. SARS-CoV-2 501Y.V2 escapes neutralization by South African COVID-19 donor plasma. Nat Med 27, 622–625, doi:10.1038/s41591-021-01285-x (2021).

15 Hoffmann, M. et al. SARS-CoV-2 variants B.1.351 and P.1 escape from neutralizing antibodies. Cell 184, 2384–2393 e2312, doi:10.1016/j.cell.2021.03.036 (2021).

16 Edara, V. V. et al. Infection and Vaccine-Induced Neutralizing-Antibody Responses to the SARS-CoV-2 B.1.617 Variants. N Engl J Med 385, 664–666, doi:10.1056/NEJMc2107799 (2021).

17 Xie, X. et al. An Infectious cDNA Clone of SARS-CoV-2. Cell Host Microbe 27, 841–848 e843, doi:10.1016/j.chom.2020.04.004 (2020).

18 Francica, J. R. et al. Vaccination with SARS-CoV-2 Spike Protein and AS03 Adjuvant Induces Rapid Anamnestic Antibodies in the Lung and Protects Against Virus Challenge in Nonhuman Primates. 2021.2003.2002.433390, doi:10.1101/2021.03.02.433390 %J bioRxiv (2021).

19 Katzelnick, L. C. et al. Viridot: An automated virus plaque (immunofocus) counter for the measurement of serological neutralizing responses with application to dengue virus. PLoS Negl Trop Dis 12, e0006862, doi:10.1371/journal.pntd.0006862 (2018).

